# Interoceptive Signals Bias Decision Making in Rhesus Macaques

**DOI:** 10.1101/2024.07.08.602563

**Authors:** Michael A. Cardenas, Ryan P. Le, Tess M. Champ, Derek O’Neill, Andrew J. Fuglevand, Katalin M. Gothard

**Author notes:** **Corresponding Author** Katalin M. Gothard. **Author Contributions** MAC: designed the experiments, collected the data, analyzed the data, and wrote the manuscript. RPL: helped train the animals, helped analyze the data. TMC: set up and implemented the first version of the experiments, trained the animals. DO: trained the animals, designed and implemented the arm restraint, helped collect the data. AJF: helped design the experiments, helped analyze the data, and wrote the manuscript. KMG: designed the experiment, oversaw data collection and analyses, and wrote the manuscript. **Competing interest statement** The authors declare no competing interests.

## Abstract

Several influential theories have proposed that interoceptive signals, sent from the body to the brain, contribute to neural processes that coordinate complex behaviors. We altered the physiological state of the body using compounds that have minimal effect on the brain and evaluated their effect on decision-making in rhesus monkeys. We used glycopyrrolate, a non-specific muscarinic (parasympathetic) antagonist, and isoproterenol, a beta-1/2 (sympathetic) agonist, to create a sympathetic-dominated state in the periphery, that was indexed by increased heart rate. Rhesus monkeys were trained on two variants of an approach-avoidance conflict task. The tasks offered a choice between enduring mildly aversive stimuli in exchange for a steady flow of rewards, or cancelling the aversive stimuli, forgoing the rewards. The delay to interrupt the aversive stimuli was used as a measure of monkeys’ tolerance for contact with a hot but not painful stimulus or airflow directed at their muzzle. Both drugs reduced tolerance for the aversive stimuli. To determine whether the drug-induced autonomic state reduced the subjective value of the reward, we tested the effects of glycopyrrolate on a food preference task. Food preference was unaltered, suggesting that the sympathetic dominated state in the periphery selectively reduces tolerance for aversive stimuli without altering reward-seeking behaviors. As the drugs used are expected to have little or no direct effect on the brain, the observed biases in decision making are likely induced by interoceptive afferents that signal to the brain the physiological state of the body.

**Significance statement:** The brain adjusts body physiology to the behavioral agenda of the organism through autonomic efferents; concomitantly interoceptive afferents carry signals that inform the brain about the physiological state of the body, closing a homeostatic regulatory loop. Persuasive theories proposed that interoceptive afferents contribute to higher cognitive functions, including emotion. Empirical evidence that these signals are sufficient to bias complex behavior has been limited by the difficulty of isolating interoceptive afferents from the rest of the homeostatic loop. Here we selectively manipulated the autonomic state of the body using drugs with limited penetrance of the brain in macaques performing decision-making tasks. Sympathetic-dominated peripheral states significantly altered decision making, suggesting that changes in interoceptive afferent signals are sufficient to bias behavior.

## Introduction

The long-standing debate regarding the role of body physiology in shaping mental/emotional experiences was ignited by the James-Lange theory, claiming that distinct physiological states precede and define emotions (James, 1884, 1890; Lange, 1885; Papez, 1937). The opposing Cannon-Bard theory emerged from the empirical failure to match distinct autonomic states to each emotion and from the observation that emotional behaviors remain intact after lesions that prevent the brain from sensing the state of the body (Bard, 1928; Bard & Rioch, 1937; Cannon, 1927, 1931; Sherrington, 1900). Following partially successful attempts to reconcile these theories though a cognitive model of emotion (Schachter & Singer, 1962), new models emerged that expanded the role of brain-body interactions into higher cognitive spheres (Adolphs & Anderson, 2018; Barrett, 2017; Bechara & Damasio, 2005; Damasio, 2012; Damasio, 1996; Feldman et al., 2024; Varela et al., 2017). The current theories rest on a more nuanced understanding of the interplay between regulatory homeostatic processes in the autonomic nervous system and social, emotional, and cognitive processes in higher-level brain circuits. Despite the well-established links between body physiology, subjective emotional experience (Critchley et al., 2005; Garfinkel et al., 2014; Gray et al., 2012), decision making (Werner & Schandry, 2024), and the sense of self (Barrett, 2017; Barrett & Simmons, 2015; Damasio, 2012), it is unclear whether changes in body physiology are *sufficient* to alter cognitive-emotional processes in the brain.

Answering this question is hindered by the closed-loop architecture of the central autonomic network that articulates to each other the ascending and descending limbs of brain-body circuits (Benarroch, 1993; Berntson & Khalsa, 2021; Ferraro et al., 2022; Quadt et al., 2022). Indeed, body physiology (e.g., heart rate) is both the *effect* of descending autonomic commands and the *cause* of ascending signals that inform the brain about the changes induced by previous commands. In turn, the ascending, interoceptive signals may precipitate further descending adjustments from central structures, thereby closing a loop. A fruitful approach to isolating the role of interoceptive afferent signals would be to alter the physiological state of the body with manipulations that selectively target the internal organs, and then measure subsequent changes in brain states and/or behaviors.

Recently, Hsueh et al., (Hsueh et al., 2023) used this approach to show that selective cardiac perturbations in mice elicits exaggerated avoidance behaviors. Likewise, tachycardia induced in humans by isoproterenol, a beta receptor agonist with minimal capacity to cross the blood-brain barrier, caused feelings of anxiety (S. Khalsa et al., 2009; Verdonk et al., 2024). The question left unanswered by these studies is whether interoceptive states are sufficient to bias more elaborate behaviors.

Elaborate evaluation of costs and benefits are required for making decisions in approach-avoidance conflict tasks (Aupperle et al., 2011; Kirlic et al., 2017; Letkiewicz et al., 2023; Vogel et al., 1971; Walters et al., 2019). We trained non-human primates to perform different versions of approach-avoidance conflict tasks and leveraged pharmacological agents with limited blood-brain barrier penetrance that altered the physiological state of the body (Ali-Melkkilä et al., 1990; Ali-Melkkilä et al., 1989; Ali-Melkkilä et al., 1990; Ali–Melkkilä et al., 1993; Olesen et al., 1978; Proakis & Harris, 1978). Such manipulations helped to isolate the effect of afferent interoceptive signals on decision making. We found that application of these peripherally restricted drugs had substantial effects on choice behavior, demonstrating that changes in body physiology can induce reliable changes in decision-making.

## Results

### Pharmacological manipulations

We identified three drugs that alter sympathetic/parasympathetic balance in the periphery (henceforth, autonomic state) with limited penetrance of the blood-brain barrier. Two of the drugs, glycopyrrolate (a non-selective muscarinic receptor antagonist) and isoproterenol (a non-selective beta-receptor agonist), shift autonomic state toward sympathetic dominance. The third drug, atenolol (a cardioselective beta receptor blocker), blunts the heart’s response to sympathetic inputs. We elected to use heart rate as a general readout of autonomic balance, although we expect isoproterenol and glycopyrrolate to act on multiple organs. In healthy adult macaques, glycopyrrolate increased heart rate relative to saline control (**Figure 1**) by 20.5 ± 6.7 (mean ± SD) beats per minute (BPM) (Wilcoxon rank-sum test p<0.01). Likewise, isoproterenol (**Figure 1**) increased heart rate by 11.3 ± 6.0 BPM (Wilcoxon rank-sum p< 0.01). Atenolol caused a large decrease in heart rate in monkey A (−26.1 ± 5.2 BPM) but had no effect in monkey S (3.7 ± 4.5 BPM).

**Figure 1:**
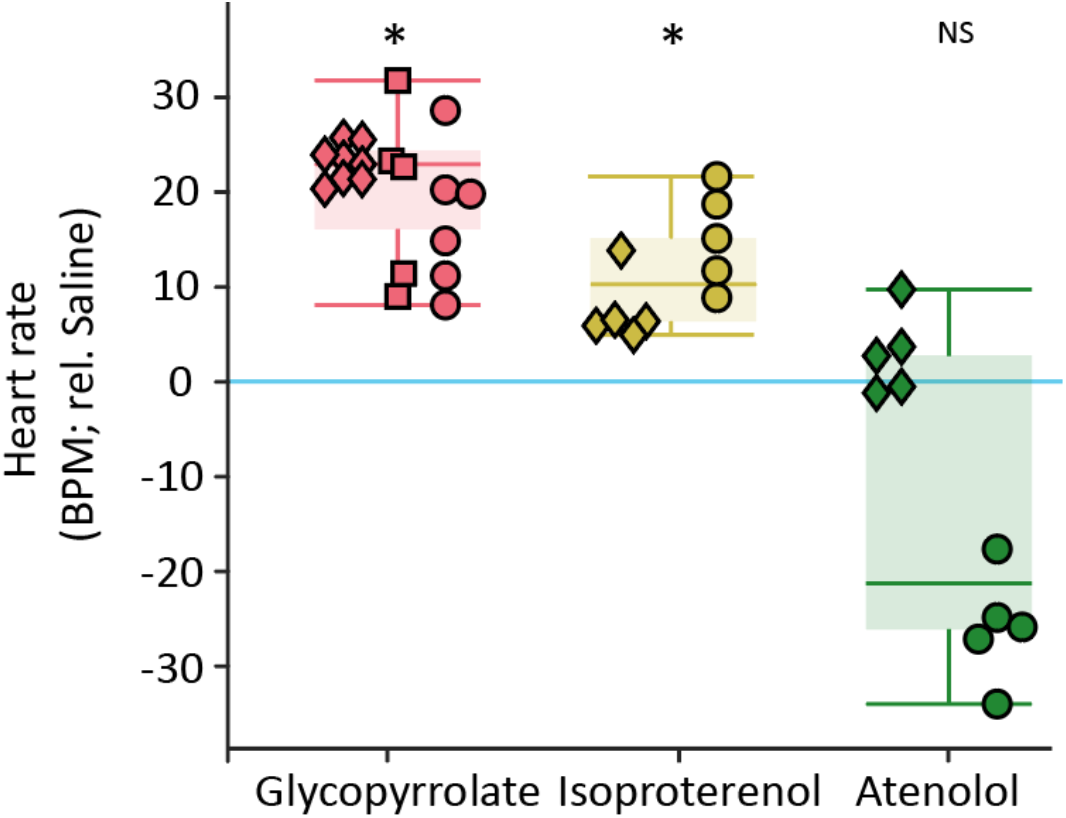
Effects of glycopyrrolate, isoproterenol and atenolol on heart rate in two male (A and S) and one female (P) adult macaques (monkey S = diamond, monkey P = squares, monkey A = circle). Glycopyrrolate (n = 19 sessions) and isoproterenol (n = 10 sessions) led to significant increases in heart rate (p < 0.01, Wilcoxon rank-sum test) compared to saline injection. Atenolol (n = 10 sessions) decreased heart rate in one monkey only (p = 0.065 Wilcoxon rank-sum test). Asterisks represent significant change in heart rate relative to saline.

### Approach-avoidance conflict task

We designed a novel approach-avoidance conflict task in which animals chose between two options: (1) endure a hot but non-painful stimulus in exchange for a steady flow of fruit juice, or (2) turn off the heat by using a switch, forgoing the juice reward. A thermode (a Peltier machine that rapidly heats and cools) was attached to a shaved region of the monkeys’ arms (**Figure 2A**). On each trial, the temperature was set either to remain at a neutral temperature of 35°C, (*no-heat trials*), or to ramp up over 1s to a predetermined hot but not painful temperature ranging between 46 and 48°C, depending on the tolerance of the monkey (*heat trials*). While the heat remained on (for a maximum of 20 s), monkeys received a 0.2 ml drop of fruit juice at a rate of 1 drop/s. At any time after the heat began to increase, monkeys could interrupt heat and juice delivery by touching the switch for 600 ms. The latency to turn off the heat served as a measure of the animal’s tolerance to the mildly aversive heat stimulus in exchange for receiving the juice reward.

**Figure 2.**
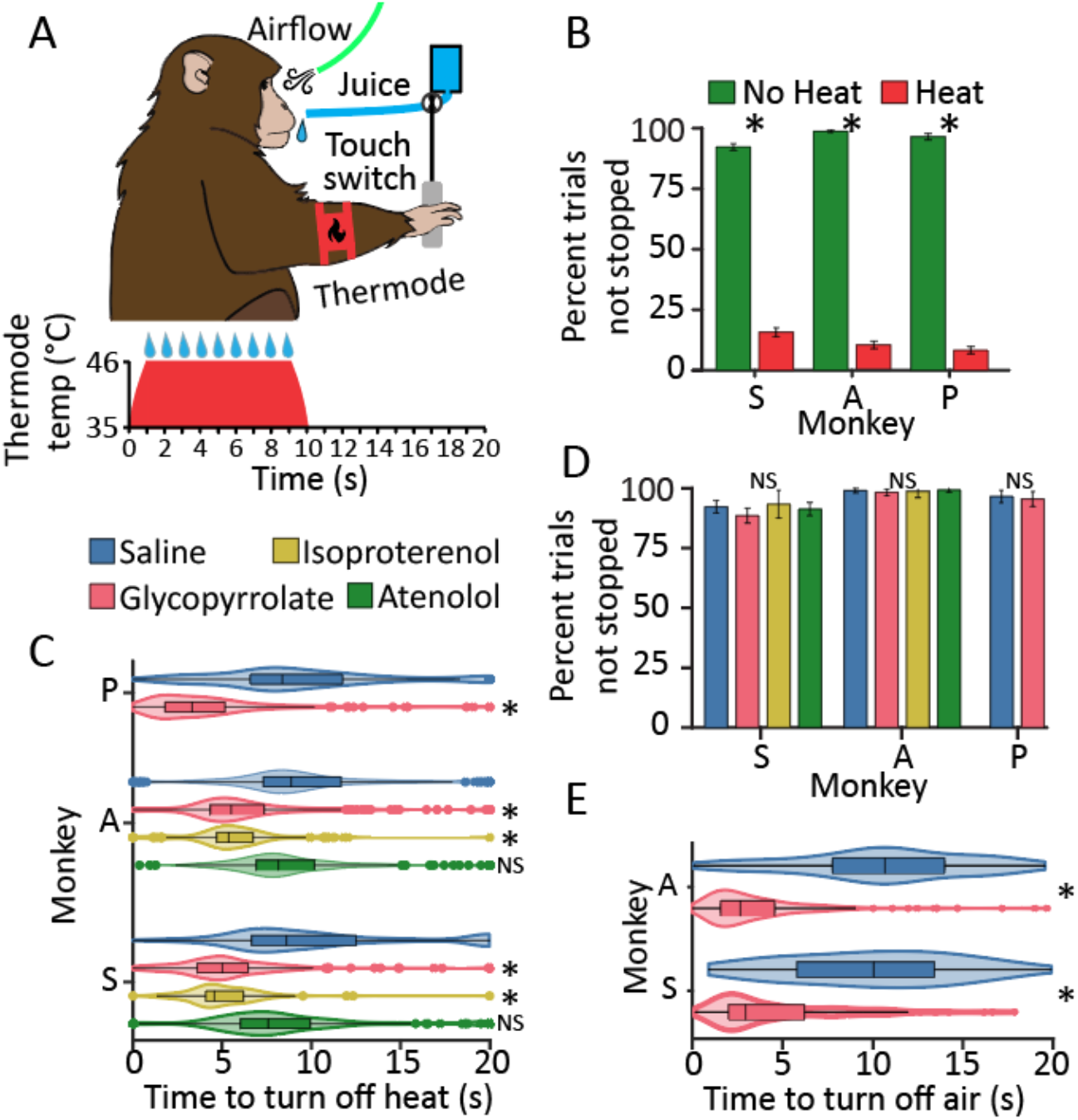
Effects of pharmacological manipulations on behavior during approach-avoidance conflict. **A**: Cartoon of the approach-avoidance conflict task with the thermode or airflow as the aversive stimulus. At the start of a trial, the thermode rapidly heated up to a predetermined temperature or the airflow was turned on and remained at a constant pressure for a maximum of 20 s. A drop of juice was delivered every second while the heat or airflow was on (red trapezoid). The monkey could interrupt the aversive stimulus by touching a switch, but this also halted juice delivery (in this example trial the monkey interrupted the aversive stimulus at ∼9 s). **B**: Proportion of trials not stopped by the monkeys during heat and no heat trials when no drugs were delivered (N=400, 400, and 200 trials for monkeys S, A, and P respectively; Chi-squared test of proportions p<0.001). **C**: Latency to turn off the heat under different drug conditions (number of trials = monkey A: saline = 400, glycopyrrolate = 400, isoproterenol =150, atenolol = 400; monkey S: saline = 400, glycopyrrolate = 400, isoproterenol = 150, atenolol = 400; monkey P: saline = 300, glycopyrrolate = 300). Asterisks represent significant change in behavior relative to saline (Wilcoxon rank-sum test p<0.001). **D**. Proportion of trials not stopped by the monkeys during the no heat trials under different drug conditions (Chi-squared test of proportions p>0.05; number of trials: monkey A: Saline= 400, glycopyrrolate = 400, isoproterenol = 75, atenolol = 400; monkey S: saline = 400, glycopyrrolate = 400, isoproterenol = 75, atenolol = 400; monkey P: saline = 200, glycopyrrolate = 200). **E**. Latency to deactivate airflow after saline or glycopyrrolate administration (200 trials each; Wilcoxon rank-sum test p<0.001).

Each experimental session consisted of a block of 15 - 50 no-heat trials and a block of 30 - 50 heat trials. The order of the blocks was randomized across sessions. As the animals were not water restricted, the number of trials in each session varied with their level of satiation. The three animals tested on this task rarely stopped the no-heat trials but stopped the majority of heat trials (Chi-Squared test of proportions, *p*<0.001 **Figure 2B**), indicating that the presence of the heat biased their behavior.

### A sympathetic-dominated state reduces tolerance for aversive stimuli

Under control (saline) conditions all monkeys tolerated the heat for an average duration of 7-9 s (**Figure 2C**, blue violin plots, overall average 8.6 ± 3.6 s). Glycopyrrolate administration shortened heat tolerance to an average latency of 5.1 ± 3.1 s (a 3.5 s or 40.7% decrease) (**Figure 2C**, pink violins; Wilcoxon rank-sum *p*<0.001; Cohen’s d: monkey A: 0.8, monkey S: 1.0, monkey P: 1.5), indicating that blocking peripheral parasympathetic receptors reduced heat tolerance. To determine if the reduced heat tolerance was the result of a sympathetic-dominated autonomic state and not the specific result of a muscarinic blockade, we replaced glycopyrrolate with isoproterenol in monkeys A and S. Isoproterenol, like glycopyrrolate, reduced the duration of heat tolerance in both monkeys to 5.3 ± 2.1 s (a 3.3 s or 38.4% decrease) (**Figure 2C** yellow violins; Wilcoxon rank-sum *p*<0.001; Cohen’s D: monkey A: 1.0 monkey S: 1.1). Thus, blocking parasympathetic muscarinic receptors or activating sympathetic beta receptors had the same effect of increasing heart rate and reducing heat tolerance. Next, we tested whether shifting sympathetic/parasympathetic balance toward parasympathetic dominance by blocking the sympathetic cardiac receptors with atenolol would have the opposite effect. While atenolol decreased heart rate in one monkey, it had no effect on approach-avoidance decisions in either subject (**Figure 2C**, green violins; Wilcoxon rank-sum p>0.05). Furthermore, as the drugs had no effect on the monkeys’ behavior on the *no heat* trials, it is unlikely that they altered reward seeking behavior or devalued the reward (**Figure 2D**; Chi-squared test of proportions compared to saline control, *p*>0.05).

To determine if the effects of isoproterenol and glycopyrrolate were specific to thermal stimuli, or generalize to other aversive stimuli as well, monkeys A and S performed an airflow version of the approach-avoidance conflict task. In this case, for each trial, monkeys either endured a continuous flow of room-temperature air directed at their muzzle for 20 s or could turn off the airflow and forgo the juice reward by touching the switch. Airflow pressure was set to a fixed value of ∼ 64 Pa (roughly equivalent to the air pressure delivered by a moderate breeze). Here too, the latency to turn off the airflow served as a measure of the animal’s tolerance to the mildly aversive stimulus. Eight experimental sessions were performed by both animals (four each for saline and glycopyrrolate) and each experimental session consisted of 50 trials. As in the case of the thermal stimuli, glycopyrrolate markedly reduced both subjects’ tolerance to continuous airflow directed at their face, from 10.3 ± 4.7 s with saline to 4.0 ± 3.4 s with glycopyrrolate, a 61.2 % drop (**Figure 2E**, blue and red violins; Wilcoxon rank-sum *p*<0.001; Cohen’s d: monkey A: 1.3; monkey S: 1.7). Collectively, these results suggest that a sympathetic-dominated autonomic state in the periphery reduces tolerance to aversive outcomes, regardless of the nature of the stimulus.

### A sympathetic-dominated state does not alter decision making for non-aversive stimuli

To determine whether sympathetic-dominated states alter decisions about rewards, we evaluated the effects of glycopyrrolate on a task where the subjects chose between appetitive stimuli only. On each trial, monkeys were offered two foods randomly selected from a pool of four possible foods. Pictures of the offered foods were presented on a screen and the monkeys made their selection by fixating for 500 ms on the picture of their preferred food. Two monkeys (monkeys C and P) performed this food preference task before and after receiving a glycopyrrolate injection (36 pre-administration and 36 post-administration trials). Each monkey participated in 8 sessions. We limited the number of trials within a session to minimize satiation effects. We hypothesized that if decision-making was generally disrupted by glycopyrrolate, the monkeys might change their food preference.

Glycopyrrolate did not alter absolute or ordinal food preference (**Figure 3**; Friedman’s test *p*>0.05). Thus, a sympathetic-dominated peripheral autonomic state has little effect on decision making regarding appetitive stimuli.

**Figure 3:**
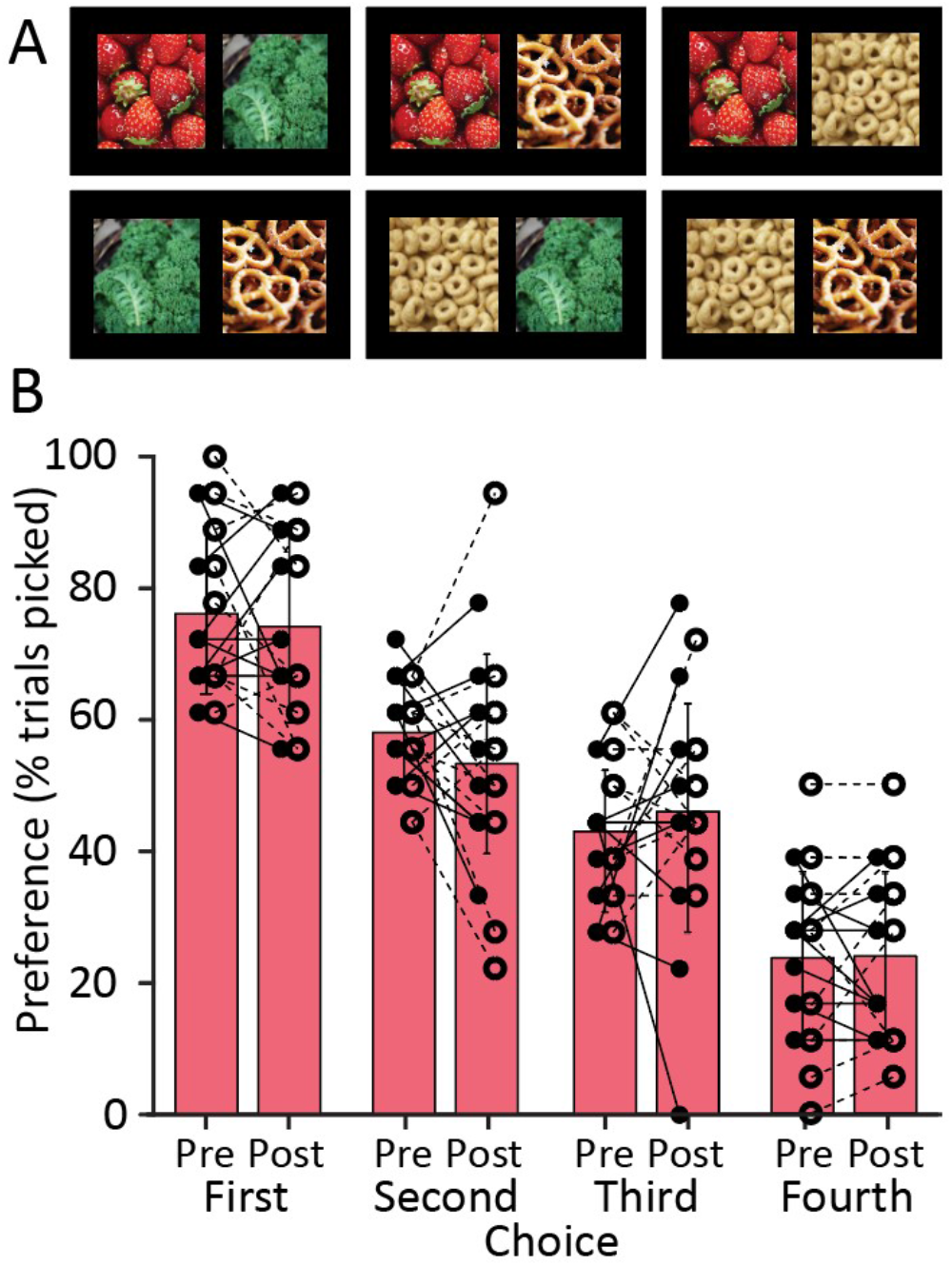
Glycopyrrolate did not change ordinal food preference. **A**. Six possible pairs of four foods used in the food preference task. All six pairs were tested. On each trial, the monkeys were shown a pair of images. The monkeys indicated which food they prefer by fixating on the image for 500 ms, after which the image of non-preferred food disappeared from the monitor and a human handler delivered the chosen food to the monkey. **B**. Food preference pre and post glycopyrrolate injection. Each pair of markers connected by a line correspond to the proportion of trials in which a food item was chosen before and after glycopyrrolate administration. Filled dots and lines are for monkey P while circles and dashed lines are for monkey C. Monkeys’ food preferences before injection were ordinally ranked with the leftmost bars indicating the most preferred food and rightmost bars the least preferred food. Error bars are SD. Glycopyrrolate administration did not modify food preference (Friedman’s test p>0.05).

## Discussion

The results presented here demonstrate that peripherally restricted manipulations of body physiology are sufficient to bias decision making in two versions of an approach-avoidance conflict task. Despite acting on different receptors, both glycopyrrolate and isoproterenol caused elevated heart rate, indicating a sympathetic-dominated autonomic state. As both glycopyrrolate and isoproterenol bind to multiple types of muscarinic and adrenergic receptors, we expect these drugs to cause widespread changes in the functional state of multiple organs. Such altered functional states are likely detected by different sets of interoceptive receptors, and the resulting signals are transmitted to homeostatic centers in the brainstem and hypothalamus via interoceptive afferents (Craig, 1995, 2002; Saper, 2002). These afferents represent the ascending limb of regulatory loops that typically elicit compensatory or corrective central mechanisms sent back to the target organs through the descending limb of these circuits. These processing loops, however, are not fully symmetrical. While the descending limbs originate in the autonomic centers of the brainstem and hypothalamus and terminate at the level of the target organs, the ascending limb connects back to the regulatory centers, and from there, extends to the amygdala, insula, and via the thalamus, to distinct areas of the cerebral cortex (Berntson & Khalsa, 2021; Chen et al., 2021; Craig, 1995, 2002, 2003, 2009; Craig et al., 1994; Saper, 2002). The biased decision making reported here is likely the result of such interoceptive afferents influencing decision-making circuits involved in approach-avoidance tasks.

A subset of neurons that receive interoceptive inputs are located in the anterior cingulate cortex and become active during decision-making tasks (Amemori & Graybiel, 2012; Fujimoto et al., 2021). The processing of interoceptive signals may generate a conscious experience, a perception of the state of the body, which is the proper meaning of the term “interoception” (Barrett & Simmons, 2015; Engelen et al., 2023; Khalsa et al., 2018). While interoceptive sensitivity and awareness have been linked to cognitive and affective states, and to a variety of clinical conditions (Critchley & Garfinkel, 2017; Garfinkel et al., 2015; Khalsa et al., 2018; S. S. Khalsa et al., 2009), interoceptive afferents may exert an influence on cognitive processes even in the absence of awareness. As macaque monkeys are capable of cardiac interoceptive sensing (Charbonneau et al., 2022), it is possible that that the behavioral effects reported here are related to the drug-induced interoceptive state that was perceived by our animals, although this was not directly tested.

Critical for interpreting the results presented here is that the drugs used had minimal penetrance through the blood brain barrier and thus no *direct* effect on the decision-making circuits in the brain. Early studies on the pharmacokinetics of glycopyrrolate indicated poor penetrating capacity across the blood–brain barrier (Proakis & Harris, 1978) or no penetration at all (Ali-Melkkilä et al., 1990; Ali-Melkkilä et al., 1989; Ali-Melkkilä et al., 1990; Ali–Melkkilä et al., 1993). While it is unlikely that glycopyrrolate crossed the blood-brain barrier, isoproterenol has 3.8% extraction in the brain after intracarotid administration (Olesen et al., 1978). This represents a 16-fold lower level compared to the lipophilic propranolol known to have central effects (Carnovale et al., 2023).

Both glycopyrrolate and isoproterenol caused sympathetic-dominated physiological states that reduced tolerance for aversive stimuli while leaving preference for appetitive stimuli intact. It is unlikely that the decision to tolerate, or not, the heat or the airflow was informed by homeostatic imperatives such as hunger or thirst (Allen et al., 2019; Livneh et al., 2020) because the food and water intake of our animals was not restricted. The most parsimonious explanation for increased avoidance is that sympathetic states resemble states of anxiety that increase avoidant behaviors (Kirlic et al., 2017). Indeed, anxiolytic drugs (diazepam) decrease avoidant behavior in rats (Vogel et al., 1971) and monkeys trained to perform a similar approach-avoidance conflict task (Amemori & Graybiel, 2012). In contrast to sympathetic-dominated states, atenolol (which prevents the heart from responding to sympathetic drive) had no effect on avoidance. While the small sample size limits interpretation, the failure of atenolol to alter behavior might be due to the absence of an interoceptive prediction error (Feldman et al., 2024; Khalsa & Feinstein, 2018). As the effects of atenolol persist for at least 24 hours (Leonetti et al., 1980), the eight consecutive days of atenolol administration may have caused allostatic adaptations that were sufficient to minimize prediction errors.

Interoceptive signals, particularly those generated on the timescale of the cardiac, respiratory, and gastric cycle, have been shown to modulate the processing of external stimuli (Critchley et al., 2005; Dunn et al., 2010; Ekman et al., 1983; Garfinkel et al., 2014; Gray et al., 2012; Gray et al., 2009; Grund et al., 2022; Park et al., 2014; Tallon-Baudry, 2023). The changes in decision making reported here, however, persisted throughout an experimental session, suggesting that interoceptive signals contributed to a sustained change in decision-making. Similarly, recent work in which perturbations of cardiac function were induced by optogenetic stimulation of the heart, led to a persistent increase in avoidance of anxiogenic stimuli (Hsueh et al., 2023).

Our study has several limitations. Although the observed behavioral effects could be attributed to the activation of interoceptive afferents by peripheral perturbations, it is also possible that peripheral perturbation engage efferent compensatory effects (e.g., tachycardia triggers compensatory parasympathetic effects). It is possible, therefore, that regulatory efferents contribute indirectly to the observed changes in decision making (for example via an efference copy that may be sent to the decision-making circuits of (Damasio, 2012). Further, to capture the direct causal relationship between changes in body physiology and decision making, we did not modify the parameters of our approach-avoidance conflict task to develop psychometric curves by presenting multiple heat or airflow levels, intermixing trial types, randomizing the reward cadence, or varying the value of the juice reward. In addition, while the one female subject tested in this study exhibited a stronger response to glycopyrrolate administration, a much larger sample size would be required to evaluate sex-related differences.

This study represents only the first step in exploring the role of interoceptive afferents in shaping cognitive processes in non-human primates. This protocol can be expanded to test the role of ascending interoceptive signals in other cognitive domains and most importantly for further exploring the neural basis of brain-body interactions. Recording neural activity from the nodes of the central autonomic network (Benarroch, 1993; Ferraro et al., 2022; Quadt et al., 2022) would be ideal candidates for follow-up studies because they are the main recipients of the interoceptive signals and have been implicated in emotion, decision-making, and social behaviors (Berntson & Khalsa, 2021; Buzsáki & Tingley, 2023; Feldman et al., 2024; Karalis & Sirota, 2022).

## Methods

### Subjects and Living Arrangements

All protocols were approved by the University of Arizona Institutional Animal Care and Use Committee and carried out in accordance with the US National Institutes of Health guidelines.

Two female (monkeys C and P, 16 and 12 years old, respectively) and two male (monkeys A and S, both 8 years old) adult rhesus monkeys (*Macaca mulatta*) weighing, on average 11.7 ± 2.1 kg were housed in standard indoor cages in temperature-controlled rooms with automatically regulated lighting (12 h light/dark cycle with lights on at 7:00 AM and off at 7:00 PM). All monkeys were housed with a cage mate of the same sex, were allowed access to water *ad libitum* in cage, and fed once daily (Tekklad 2050) in accordance with veterinary directions around 3 pm each day. Data collection of these experiments took place typically between 9 and 11 am the next day. Plastic toys, mirrors, and other objects were provided in-cage for enrichment. Nightly enrichment including foraging boxes, vegetables, ice cubes, etc., was provided alongside their standard diet. Monkeys A and S were cage mates, as were monkeys C and P. Monkeys A, S, and P were trained on the thermal approach-avoidance conflict task, monkeys A and S were trained on the airflow approach-avoidance conflict task, and monkeys P and C were trained on the food preference task.

### Thermal approach-avoidance conflict task

Monkeys were acclimated to custom-made primate chairs and trained to place their arms in custom made arm restraints built from PVC tubes attached to a rail system inside the chair. The restraint for the right arm had an opening on its left side for the monkey to move their wrist freely and interact with a switch that was mounted on the arm restraint. The left arm restraint had an opening that snuggly fit the 30 × 30 mm head of a Medoc TSA-II Thermode (Ramat Yishay, Israel). Elastic Velcro straps were attached to keep the thermode head in contact with the skin of the dorsal proximal forearm. Care was taken not to put excessive pressure on the thermode head that might restrict blood flow or numb the skin.

Automated stimulus and juice delivery was coordinated by a custom program on the platform of the NIMH MonkeyLogic scripts (Hwang et al., 2019). The thermal stimulus was delivered using a MEDOC TSA-2 thermal sensory device. This is a Peltier device approved by the FDA for human use. Custom scripts were written to allow MonkeyLogic to interface with the MEDOC software package to control the heat in accordance with MEDOC external control documentation (TSA-2 manual appendix K). For reproducibility, we used the external control mode and created custom versions of the *TSA – 2 “42 ramp and hold”* demo program. The baseline was set to 35°C and the target heat was different for each monkey (heat ranging from 46 to 48°C). The ramp time from baseline to target was set to 1s. The switch was a battery powered PCB gadgets CP100 Capacitive Switch with the sensor connected to a metal bar and the output signal was sampled by MonkeyLogic and data acquisition systems.

After being trained to tolerate arm restraints, monkeys learned to hold the metal bar for a solid food reward when cued. They then learned to activate the switch to turn off the heat when set to 46°C by cuing them after the heat had turned on. Next, we trained the monkeys to associate touching the bar with stopping juice delivery in the absence of heat. When the monkey started to avoid touching the bar while juice was being delivered, they moved to the full task by combining the heat and juice trials together.

Once trained on the task, we adjusted heat tolerance for each animal by gradually increasing the heat delivered over 10-20 trials/heat level and assessing stopping behavior, with the temperature not exceeding 48°C. Once an animal stopped the heat on average between 7 and 10 seconds, we did not increase the heat further. We then assessed the stopping behavior on the heat for three days (50 trials/day) before beginning experimental manipulation.

Once a consistent stopping behavior for a given temperature was established, the experiments were initiated. On each trial, the heat ramped to the pre-determined temperature without a cue. Monkeys received juice at a rate of 1 drop/s beginning at the start of the ramp until the switch was touched for 600 ms, or 20s had elapsed.

Intertrial interval was 10s. Typically, ∼20-45 minutes was needed for both heat and no-heat blocks of trials with a ∼1–2-minute break between blocks.

Although we did not directly test satiety (e.g., glycemia), we found that our monkeys stopped working around 150 trials. Each drop was 0.2 ml, thus the maximum volume a juice a monkey could receive on a single 20s duration trial (20 drops) was 4 ml. If the monkey never stopped the heat, they would receive 400 ml of juice in a session of 100 trials. This hypothetical maximum was never reached. The monkeys typically consumed between 150 and 300 ml per day that amounted to 100-200 Kcal/day.

### Airflow approach-avoidance conflict task

Monkeys A and S, who had already been trained on the thermal approach-avoidance conflict task, were arm restrained as in the thermal version of the task so that they could operate the touch switch. Airflow was delivered to the face through lock line hose (Crist Instruments) placed about 12 cm from the muzzle by a compressed air system containing a solenoid. The pressure applied at the muzzle was estimated to be ∼ 64 Pa by directing the airflow at a 2-cm diameter, shallow cup mounted on a force transducer approximately the same distance away from the hose opening as that of the monkey skin receiving the airflow. The force detected was ∼ 0.02 N, yielding a pressure estimate = 0.02 N / 0.000314 m^2^ = 64 N/m^2^ or 64 Pa. Airflow was automatically turned on in conjunction with juice delivery by a custom script written on the platform of NIMH MonkeyLogic. The task parameters were otherwise identical to that of the thermal approach-avoidance conflict task.

### Heart rate measurements

Raw electrocardiogram (ECG) was collected via three 2.5 × 1 cm self-adhesive electrodes attached to a shaved region on the monkey’s back and optimized with conductive gel and amplified through a Grass amplifier. ECG was sampled at 1000 Hz using either a Spike2 (Cambridge Electronics Design) or Plexon Omniplex recording system. Raw signals were filtered using a 3 – 35 Hz bandpass filter. The timing of R-waves was identified using template matching or cluster analysis from which instantaneous heart rate (BPM) was determined.

### Pharmacological manipulations

Saline, isoproterenol (0.0001mg/kg), and glycopyrrolate (0.008mg/kg) were administered by subcutaneous injection to a shaved and cleaned (with isopropyl alcohol) region on the back of the monkey. All monkeys were trained to tolerate the injection. Atenolol (6-7mg/kg) was delivered orally as a crushed powder in a peanut butter filled tortilla. Monkeys P did not accept any food that was mixed with the Atenolol powder, likely due to its bitter taste, and therefore did not participate in the atenolol experiments. All drugs were approved by the University of Arizona’s Institutional Animal Care and Use Committee and the doses were established in consultation with the veterinary team. Atenolol and glycopyrrolate were procured from MWI Animal Health (Boise, ID) and isoproterenol was generously provided by Dr. William Stauffer at the University of Pittsburgh.

In pilot experiments with glycopyrrolate, we observed changes in heart rate ∼20 minutes after subcutaneous administration. Previous studies showed that clearance of glycopyrrolate takes 3-8 hours (Ali-Melkkilä et al., 1989; Rautakorpi et al., 1994). Thus, after monkeys received subcutaneous injections of either glycopyrrolate or saline, they were allowed to rest in the chair for 30 minutes. Heart rate was not recorded during this 30-minute period. At the end of the 30-minute wait, the monkey was placed in arm restraints, ECG pads were attached to their back, and ECG recording was initiated just before they began working on the task. Atenolol was administered 30 minutes before starting the task, just as with saline and glycopyrrolate (see Supplemental Figure 1). Heart rate was recorded while the monkeys performed the task. Average heart rate from each recording session was calculated from the inter-trial intervals of the no heat block of the experiment in monkeys A and P. In monkey S, we recorded heart rate on separate experimental sessions due to difficulties with obtaining stable ECG signal while he was engaged with the task. Excessive movement, fidgeting and scratching interferes with ECG recordings in some monkeys performing behavioral tasks (Mitz et al., 2017).

Unlike glycopyrrolate or atenolol, isoproterenol is rapidly cleared (Procaccini et al., 2019), therefore, the animals received a dose before each block. Behavioral testing was initiated soon after the isoproterenol injection.

The traces in Supplemental Figure 1 show example heart rates from separate sessions after different drug (or saline) administrations in one monkey. The “task performance” bar schematically indicates approximately when the animal was involved in the tasks. Across sessions the order of no-heat and heat trials were randomized.

### Statistical analysis

Statistical analysis was done using MATLAB version 2023a. Non-parametric tests were performed across all data. Heart rate recordings were excluded if signal was lost in at least 30% of trials. This could happen as a result of excessing movement that caused contamination with EMG and/or detachment of the adhesive ECG pads.

## Acknowledgments

We acknowledge Drs. Frank Porreca and Todd Vanderah for their expertise in identifying the pharmacological agents, Drs. Anne Martin and Clayton Moser for their input on experimental design, Nathalie Sotello for her assistance in data collection and troubleshooting equipment, Dr. Tapas Arakeri for his assistance in interfacing the MEDOC thermode control software with MonkeyLogic, and Dr. William Stauffer for providing the isoproterenol. This work was supported by NIH Grants R01MH121009-02 (to A.J.F. and K.M.G.) and 1F99NS141253-01 (to M.A.C.).

---

**FIGURE S1:**
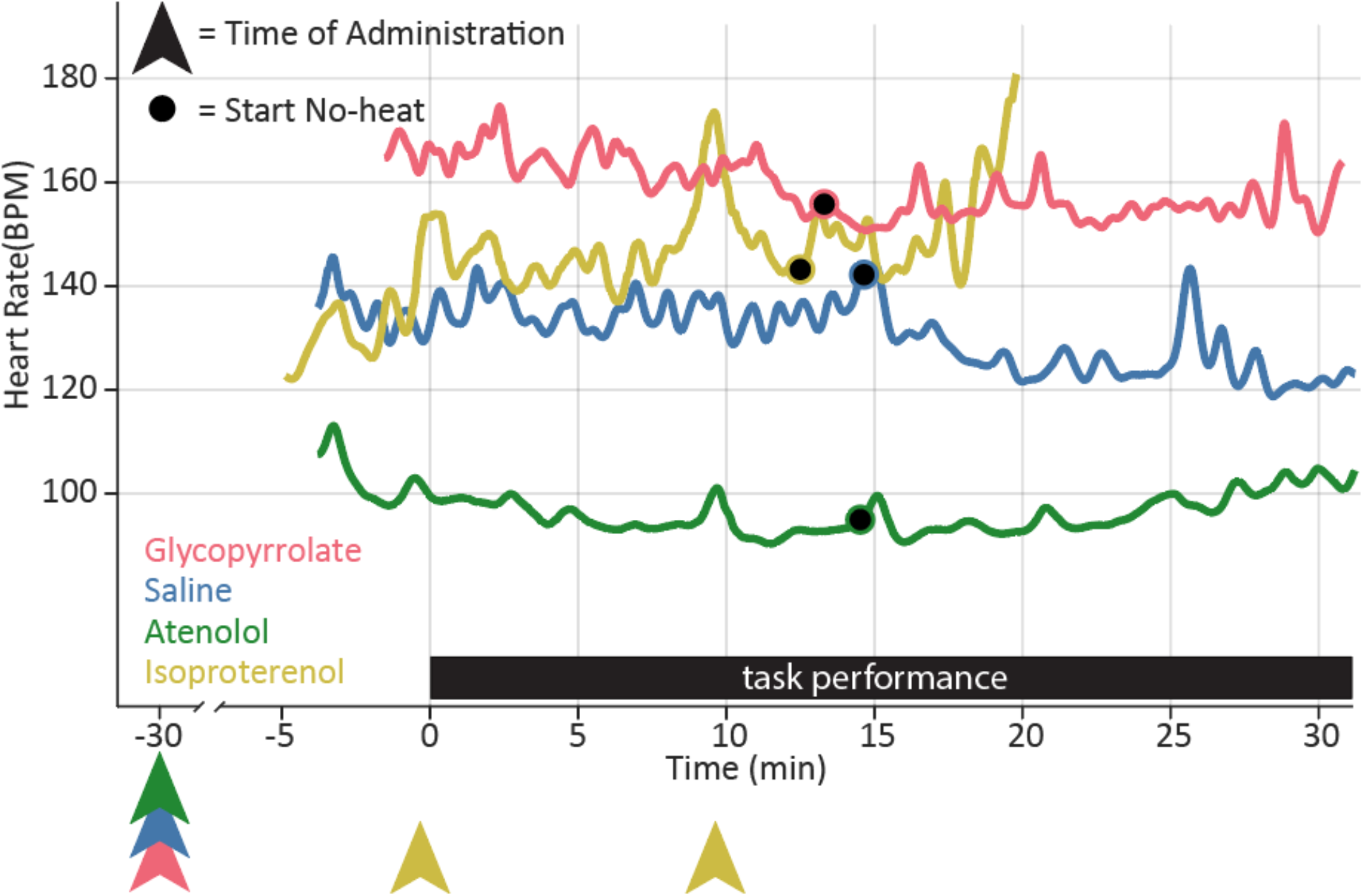
Example traces of heart rate across experimental sessions with all 3 drugs plus saline from monkey A. Instantaneous heart rate is smoothed with a 120 s gaussian filter. Arrowheads below the x axis indicate time of drug administration. The task performance bar (in black) schematically indicates the approximate time when the tasks were performed. While in some sessions, trials with no heat preceded those involving heat, in these sessions heat trials preceded no-heat trials. The black dot denotes the start of the no-heat block. Average heart rate was measured during the inter-trial intervals across the duration of the no heat trial.

