## Supplemental figure 1 for "Interoceptive Signals Bias Decision Making in Rhesus Macaques"

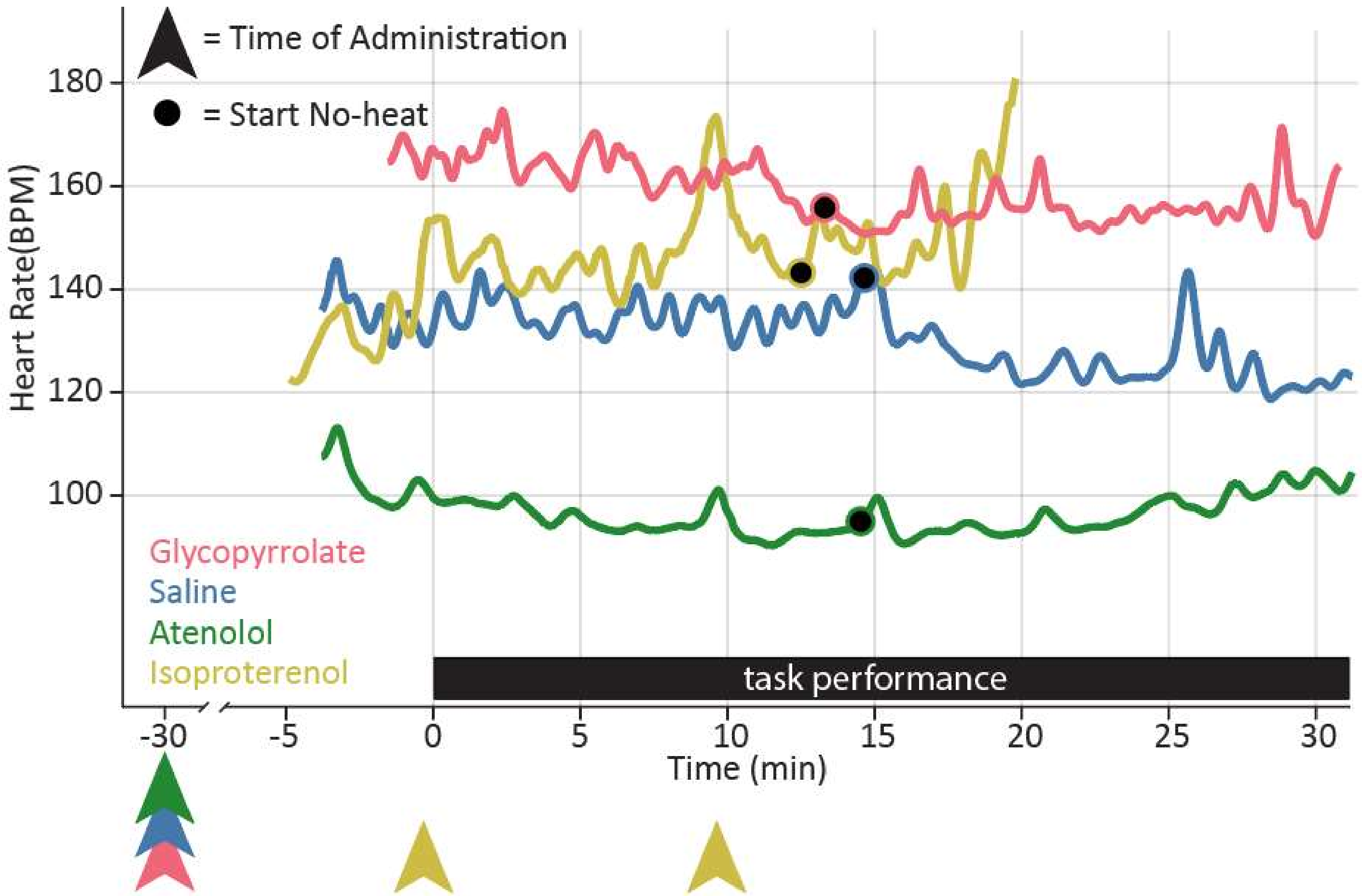


***FIGURE S1***: *Example traces of heart rate across experimental sessions with all 3 drugs plus saline from monkey A. Instantaneous heart rate is smoothed with a 120 s gaussian filter. Arrowheads below the x axis indicate time of drug administration. The task performance bar (in black) schematically indicates the approximate time when the tasks were performed. While in some sessions, trials with no heat preceded those involving heat, in these sessions heat trials preceded no-heat trials. The black dot denotes the start of the no-heat block. Average heart rate was measured during the inter-trial intervals across the duration of the no heat trial.*
